# A high-resolution single-particle cryo-EM hydrated structure of *Streptococcus pyogenes* enolase offers insights into its function as a plasminogen receptor

**DOI:** 10.1101/2022.12.06.518574

**Authors:** Sheiny Tjia-Fleck, Bradley M. Readnour, Yetunde A. Ayinuola, Francis J. Castellino

## Abstract

Cellular plasminogen (Pg) receptors (PgR) are utilized to recruit Pg, stimulate its activation to the serine protease, plasmin (Pm), and sterically protect the generated Pm from inactivation by natural host inhibitors. The net result is that cells contain a stable proteolytic surface used for biological mechanisms involved in cell migration. One such PgR is the moonlighting enzyme, enolase, some of which leaves the cytoplasm and resides at the cell surface to potentially function as a PgR. Since microbes employ conscription of host Pg by PgRs as one virulence mechanism, we explored the structural basis of the ability of *Streptococcus pyogenes* enolase (Sen) to function in this regard. Employing single-particle cryo-electron microscopy (cryo-EM), recombinant Sen from *S. pyogenes* was modeled at 2.6 Å as a stable symmetrical homooctamer displaying point group 422 (D4) symmetry, with a monomeric subunit molecular weight of ~49 kDa. Subunit-subunit interactions showed four major and four minor interfaces in the octamer. Binding sites for hPg were previously proposed to include the COOH-terminal K^434,435^ residues of Sen, but in native Sen these residues are buried within the minor interfaces of the octamer and do not function as a Pg binding epitope. Whereas Sen and hPg do not interact in solution, when Sen is bound to a surface, hPg interacts with Sen independently of K^434,434^. We propose that the octameric structure of Sen is important to its ability to interact with hPg, but disruption of its overall octameric conformation without dissociation of the octamer exposes neoepitopes for hPg binding.

## INTRODUCTION

Plasminogen receptors (PgR) are found on a variety of cells and represent an important source of cell surface proteolytic activity used by cells to migrate, disseminate, and invade (1). In the case of bacteria, such as Group A *Streptococcus pyogenes* (GAS), which is the focus of this investigation, the binding of human plasminogen (hPg) to the bacterial surface is a major virulence determinant (2). This important interaction occurs either by direct binding of hPg to its major receptors, *viz*., some subtypes of surface M-proteins (3) and/or α-enolase (Sen) (4). In these cases, activation of the bound plasminogen (Pg) by the GAS secreted endogenous activator, streptokinase (SK) (5), and/or by host-derived Pg activators (6), is highly stimulated. The resulting bound potent serine protease, plasmin (hPm), is sterically protected from inactivation by natural inhibitors (7–10), thereby allowing generation of a stable cellular proteolytic surface.

The ultimate function of the Pg activation system is to provide cell-bound hPm that can function in various pathophysiological processes, such as tissue remodeling and fibrin surveillance, as well as migratory events associated with inflammation, bacterial invasion, and cancer metastasis (11). The ubiquitous presence of PgRs on cells is consistent with its multiplicity of functions. Our specific current interests involve the role of the Pg activation system in bacterial pathogenesis. In bacterial cells, two of the more important direct binding PgRs are the GAS surface M-protein characteristic of the GAS strain that is used to serotype the strain and a group of glycolytic moonlighting enzymes, the most studied of which is cell surface enolase (Eno). There are approximately 250 strains of GAS with serotypically-distinct M-proteins, grouped in five subclasses, Patterns A-E (12). Only skin-trophic Pattern D GAS strains, a predominant Pattern type, directly and tightly interact with Pg *via* their M-proteins (PAMs) (5, 13). Some of the characteristic M-proteins of Pattern A-C GAS strains directly bind to fibrinogen, which in-turn interacts with hPg and hPm, thus providing fibrinogen as another potential cell surface PgR (14, 15).

Human Pg (hPg) is a 791-amino acid residue single-chain multi-modular zymogen that is activated by cleavage of a single peptide bond at R^561-^V^562^, resulting in the disulfide-linked two-chain protease, hPm. The COOH-terminal derived light chain is homologous to trypsin-like serine proteases and is responsible for the proteolytic activity of hPm. The amino terminal-derived heavy chain contains five ~80 amino acid triple disulfide bonded kringle (K) domains, numbered consecutively and spaced by linker regions. With the exception of K3, each of the kringle modules is independently capable of interaction with lysine (16). This property of the kringle domains is generally believed to be responsible for the binding of hPg and hPm to its receptors on cells. These PgRs either possess a carboxy-terminal lysine residue (4) or have side-chain arrangements with appropriately spaced lysine and aspartate/glutamate side-chains that provide through-space lysine isosteres (17). PAM-type M-proteins utilize this latter mechanism to tightly bind to the K2 module of hPg/hPm. Other receptors, such as Eno, are thought to utilize carboxy-terminal lysine residues as well as isosteric lysines to interact with the lysine binding sites (LBS) of K1, K4, and/or K5, in order to stabilize hPg on many types of cells, including GAS (18).

Eukaryotic enolases generally exist as homodimers and prokaryotic enolases are, in many cases, homohexamers or homooctamers. In order to understand the forces that stabilize the multimeric enolases, and the epitopes that are responsible for hPg/hPm binding in these higher order complexes, high resolution structures of these proteins are required. In this communication, we provide a single-particle 2.6Å 3D-structure of homooctameric enolase from GAS in (vitreous) ice and mine this structure for features that stabilize the octameric form of this protein and that identify the possible locations of hPg binding epitopes. Such knowledge will be of great value in modeling interactions of enolase with its ligands, thereby providing means to inhibit these host-microbial interactions that take place under pathological conditions.

## MATERIALS AND METHODS

### Expression and Purification of Sen

The nucleotide sequence of *Streptococcus pyogenes enolase (sen)* (GenBank: AMY97107.1) from Group A *Streptococcus pyogenes* (GAS) isolate AP53 (19) was used to generate recombinant (r) Sen. For this purpose, PCR amplification of the open reading frame (ORF) encoding the *sen* gene was accomplished using primers P1 (SEN-WT-F; *Nde1*) and P2 (SEN-WT-RP; *BamH1*) (**Supplemental Table 1S**). The resulting amplicon was subjected to DNA sequencing and encoded M^1^ (ATG) to K^435^ (AAA) of *sen*. The *sen* cDNA was ligated into plasmid blunt II Topo (Invitrogen) and then transformed into TOP10 competent *Escherichia coli* cells. The *sen* gene was digested at its 5’ and 3’ *NdeI* and *BamHI* sites that were engineered into *sen* during the construction. The gene was then inserted into *E. coli* expression plasmid *pET-15b* (Novagen), using these same sites, thus providing *pET15b-sen*. The plasmid contained the ampicillin resistance gene and a NH2-terminal (His)_6_-tag.

Plasmid *pET15b-sen* was transformed into *E. coli* BL-21 (DE3) cells and grown in Luria-Bertani broth or on Luria-Bertani agar plates with ampicillin (50 mg/mL). Single bacterial colonies were grown overnight in 50 mL media and transferred to 1 L media at 37° C until an OD_600nm_ of ~0.6 was reached. Expression of Sen protein was induced with 1 mM isopropyl-ß-D-thiogalactopyranoside (IPTG) for 5 hr at 30° C. The cells were then collected by centrifugation, lysed, and Sen was purified using Ni^+^-agarose affinity chromatography. The full protocol for Sen purification has been published previously (20). The protein purity was evaluated using Coomassie blue staining with SDS-PAGE gels and Western analysis using in-house rabbit-anti Sen.

### Generation of Sen Variants

Point mutations at desired locations of the *sen* gene were introduced using a series of primers (**Supplemental Table 1S)** to amplify the ORF of *sen*. P1 (SEN-WT-FP; *Nde1*) in conjunction with P5 (SEN-434,435-RP) was used to change the last two codons encoding K^434^ and K^435^ into A^434^ and A^435^ (Sen[K^434,435^A]). Point mutations at K^252^ and K^255^ (Sen[K^252,255^A] were generated by making two different *sen* gene fragments using primers P1 and P3 (SEN-252,255-RP), and P4 (SEN-252,255-FP) and P2 (**Supplemental Table 1S)**. The PCR-amplified DNA fragments obtained from each pair of primers were further amplified using P1 and P2 to obtain Sen[K^252,255^A]. The *sen* gene fragments amplified using P1 and P5 were used to generate Sen[K^252,255,434,435^A].

Expression and purification of the Sen variants were accomplished as above for WT-Sen after verification of the mutagenic changes through nucleotide sequencing.

### Sen Enzymatic Activity

The activities of Sen and Sen variants were measured at room temperature in 50 mM NaH2PO4/100 mM NaCl/10 mM MgCl2/3 mM 2-phosphoglycerate, pH 7.4, in a final volume of 200 μL (9). The conversion of 2-phosphoglycerate (2-PG) to phosphoenolpyruvate (PEP) was determined by assay of PEP formation at A_240nm_ (21). Measurements were recorded every 5 sec for a total of 3 min.

### Analytical Ultracentrifugation (AUC)

The sedimentation velocities (SV) and molecular weights (M.Wt.) of Sen and Sen variants were determined by AUC in 50 mM Na^+^ phosphate/0.1 M NaCl, pH 7.4, or 50 mM NaHCO3, pH 9.6, using absorbance optics at 280 nm. SV runs were performed at 30,000 rpm with the An-60 Ti rotor and double-channel centerpiece cells at 20° C. Radial scans were recorded at 280 nm every 3 min for 16 hr at three different protein dilutions (0.8, 0.4, and 0.2 mg/mL) and analyzed by Sedfit using a continuous *c(s)* distribution model (22). The viscosities and densities of the buffer were obtained using Sednterp (http://www.rasmb.org/sednterp). Partial specific volumes of the proteins were calculated from their amino acid sequences with a subroutine in sednterp. The molecular weights and standard sedimentation coefficients (S°_20,w_) were obtained by fitting to Sedfit (https://spsrch.cit.nih.gov/). The normalized sedimentation coefficient distribution c(s*) vs* S°_20_,w plots were generated using public domain software (https://www.utsouthwestern.edu/labs/mbr.software).

### Binding of Human Plasminogen (hPg) to Immobilized Sen by ELISA

The wells of a microtiter plate were coated overnight with the Sen variants (100 μg) in PBS, pH 7.4, after which the wells were washed 3x with PBS, then incubated with hPg (0 −1.6 μM), followed by incubation with HRP-conjugated mouse anti-hPg IgG. The color was developed with 3, 3’, 5, 5’-tetramethylbenzidine and reaction stopped during the linear phase (3 min) of color development by addition of 2 N H_2_SO_4_. The A_450nm_ was determined and plotted against the hPg concentration. Graph Pad Prism software was used to plot the data and obtain the C_50_ values by-best fit of the experimental data.

### Binding of Sen Variants to Dioleoylphosphatidyglycerol (DOPG) Vesicles as Analyzed by Flow Cytometry (FCA)

The interaction of Sen variants with DOPG phospholipid (PL) vesicles was investigated essentially as described earlier (20). In summary, 0.1 mM PL vesicles pre-blocked with 2.5% BSA/PBS, pH 7.4, for 2 hr, were incubated with 0 or 10 μM WT-Sen or Sen variants for 1 hr at ambient temperature. The PL suspension was washed and centrifuged at 16,000 x g for 10 min to remove unreacted Sen. The sample was then resuspended in PBS. The formation of the PL/Sen complex of each Sen variant was then assessed by incubation of the PL with polyclonal rabbit anti-Sen for 1 hr, followed by a 30 min incubation with Alexa Fluor 488-chicken-anti-rabbit IgG (Invitrogen). Fluorescence data were acquired using a BD FACSAria III (BD Biosciences) at a flow rate of 10 μl/min with 10,000 events per acquisition. PL particles were detected by gating on fluorescence (FITC-A) and side scatter (SCC-H) using a logarithmic scale. The data were analyzed and represented in dot plots and histograms using FCS Express Version 7 software (De Novo Software, Los Angeles, CA). The FITC-positive gate was selected relative to the blank experiment by gating on PL particles with FITC intensity above those of the blank assay. The percentage of PL particles positive for FITC and the degree of positivity (median fluorescence intensity-MFI), both of which measure the ability of Sen to interact with the PL, were recorded. The data were plotted using GraphPad Prism software.

### Binding of hPg to Sen/DOPG as Measured by FCA

Replicate samples of the DOPG and Sen/DOPG complexes prepared above were incubated with hPg (0 or 0.8 μM). The binding of hPg to DOPG and Sen/DOPG was determined by FCA as described above except that mouse-anti-hPg and Alexa Fluor 488-donkey-anti-mouse IgG were used for the analysis.

### Enhancement of Tissue-type Plasminogen Activator (tPA)-catalyzed hPg Activation by Sen Variants

PL complexes of WT-Sen, Sen[K^252,255^A], Sen[K^434,435^A], and Sen[K^252,255,434,435^A], were prepared as described above. An aliquot of each PL-Sen variant complex was added to wells of a 96-well microtiter plate and incubated with 0.4 μM hPg for 1 hr at ambient temperature. Activation of hPg was initiated by the addition of a mixture of 0.25 mM of the chromogenic hPm substrate, H-D-Val-l-Leu-l-Lys-p nitroanilide (S2251), Chromogenix) and tPA (3.5 nM), diluting the final concentration of hPg to 0.2 μM. Hydrolysis of S2251 by the generated hPm was continuously monitored in a spectrophotometer by recording the A_405nm_ for 120 min. Slopes of the linear regions of plots of A_405 nm_ versus t were obtained using GraphPad Prism 9.0 and rates of hPm formation were calculated as described earlier (23).

### Sample Preparation for Cryo-EM Analysis of Wild-type (WT) Sen

Sen (3 μL of a 0.5 μM solution in 50 mM sodium phosphate buffer, pH 7.4) was pipetted onto glow-discharged CF-1.2/1.3 Holey carbon 300 mesh grids (Protochips, Morrisville, NC). The grids were blotted for 4 sec at a blotting force of 4 at 4° C/100% humidity and plunge-frozen in liquid ethane for high-speed vitrification using the FEI Vitrobot MK IV (ThermoFisher). The grids containing the sample were then loaded into the FEI Titan Krios transmission electron microscope. Data were collected over 24 hr at 300 kV using the Gatan (Pleasanton, CA) K3 direct electron detector with a Gatan Quantum GIF energy filter in super-resolution mode. Images were processed using SerialEM (Boulder Labs, Boulder, CO) software at a nominal magnification of 105,000x, resulting in a calibrated pixel size 0.437 Å. The total exposure dose was 61.37 e^−^/Å^2^, with a spherical aberration coefficient of 2.7 mm.

### Image Processing and 3D Mapping of Cryo-EM Images

All image processing was performed on CryoSPARC v3.3.1 (24) (http://www.cryosparc.com). Images were aligned and dose-weighted, and potential drift was removed by Patch Motion correction (25). The reference for focus correction was determined during the initial run. Contrast transfer function (CTF) was estimated for the dose-weighted micrographs using patch CTF estimation (26). Patch CTF extraction was performed to remove any micrographs that fell too far outside of the accepted focus or in which thick crystalline ice prevented data collection.

### Model Building and Refinement

An initial low-resolution model was generated with Phenix 1.20 Map to Model using the known Sen sequence as a template. The model was real-space refined to fit the cryo-EM map with the highest level of accuracy. The model was then imported into USCF-ChimeraX 1.2.5 for further refinement. Individual amino acids were replaced based on the known sequence. The section of the structure that was missing was input manually using ChimeraX and the missing sections of the structure were added. Once the amino acids were established, the 2° structure was corrected using the existing crystal structure (PDB:1W6T) of enolase from *Streptococcus pneumoniae* as a reference. The refined model was then imported back into Phenix to correct amino acid numbering, then again real space refined to best-fit the 3D map. The map was then measured for rotamer and Ramachandran outliers. These outliers were iteratively removed and the model re-refined until amino acids were optimally placed within the cryo-EM map.

### Resolution Determination and Structure Validation

Map and model were validated using the Phenix Cryo-EM validation tool. T he full finalized map and corresponding half maps were analyzed using Fourier Shell Correlation (FSC) at a 0.143 threshold to determine the final resolution of the WT-Sen map to be 2.63Å. The fit between the cryo-EM map and the resulting model were also determined using FSC plotting at a 0.5 threshold giving a final fit of 2.8Å between the two.

Additional results of the Phenix validation are provided in **(Supplemental Table 2S)**.

### Statistical Analyses

Statistical analyses were performed using GraphPad Prism 9.0. Error values were expressed as mean ± SD of at least four experimental replicates. Pairwise comparison was accomplished using ordinary one-way analysis of variance (ANOVA) with Dunnett’s T3 multiple comparisons test. Probability value (p-value) considered significant was set at p < 0.05.

## RESULTS

### Homogeneity and Molecular Weights of Recombinant Sen

Expression in *E. coli* and purification of Sen and Sen variants using Ni^+^-agarose gave final yields of 65-130 mg per liter of culture fluid at purities >95%. The entire process has been previously described in detail (20). SDS gels showed distinct protein bands of Sen which migrated relative to the protein markers with approximate molecular weights of ~50 kDa **(Figure 1A)**, consistent with the expected molecular weight of monomeric Sen that was calculated from the amino acid sequences of the constructs. The minor differences in mobilities of some of the variants are likely due to small changes in the conformations of the final SDS-induced structures. Western analysis of the Sen bands (**Figure 1B**) revealed that all the Sen variants are equally reactive towards a rabbit antibody raised against WT-Sen (20).

**Figure 1.**
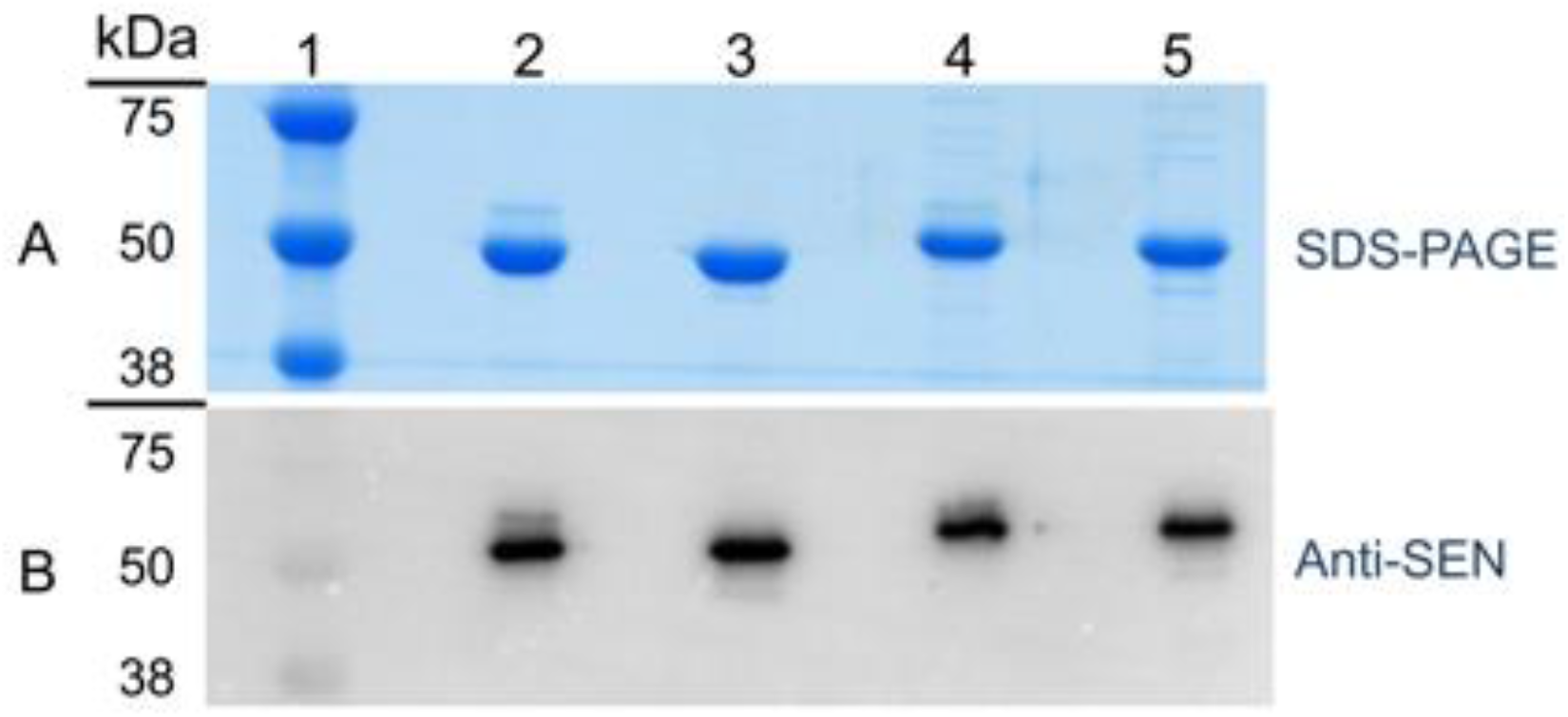
Gel analysis of purified recombinant Sen. **(A)** Coomassie Blue stained 12% SDS PAGE gels of purified Sen and its variants. Lane 1: standard molecular weight ladder; lane 2, WT-Sen; lane 3, Sen[K^252,252^A]; lane 4, Sen[K^434,434^A]; lane 5, Sen[K^252,255,434,435^A]. The Sen sample puritities are greater than 95% as confirmed by AUC; **(B)** Western analysis. As in **(A)** except that the gel was developed using rabbit-anti Sen and HRP-anti-rabbit IgG.

### Characterization of Sen Variants

WT-Sen from GAS cells has been previously characterized by AUC and SEC-MALS as a very stable homooctamer (20). At pH 7.4, all Sen variants displayed sedimentation coefficients (S°_20,w_) between 14.7 - 15.2 ± 0.2 S, which are very similar to that of WT-Sen of 14.5 ± 0.2 S (**Figure 2 A**). The small differences in the S°_20,w_ values found among the variants may be due to minor conformational changes as a result of mutagenesis. Since these latter experiments were carried out at very low concentrations of each recombinant Sen (<0.2 mg/mL), it is clear that all variants are at least 95% in their octameric structures at pH 7.4 and do not readily dissociate at low protein concentrations. In other studies, solid state binding experiments were carried out to investigate residues critical for hPg binding by coating Sen on microtiter plates at pH 9.6 (9, 27). Therefore, we further characterized the oligomeric state of Sen at pH 9.6 in order to examine the effect of pH on the octameric state of Sen which might influence hPg binding (**Figure 2B**). The data show that, similar to the observation at pH 7.4, > 95% WT-Sen and Sen[K^252,255^A] exist as octamers at pH 9.6. However, variants in which K^434^ and K^435^ were replaced by alanine showed two populations, 16-25% of which had S°_20,w_ values of 3.3 ± 0.1S, consistent with Sen monomers. These data suggest that the C-terminal lysine residues play a role in stabilizing the octameric quaternary structure of Sen and their mutagenesis to larger more hydrophobic residues, such as Leu, or their removal (9), diminishes the stability of octameric Sen.

**Figure 2.**
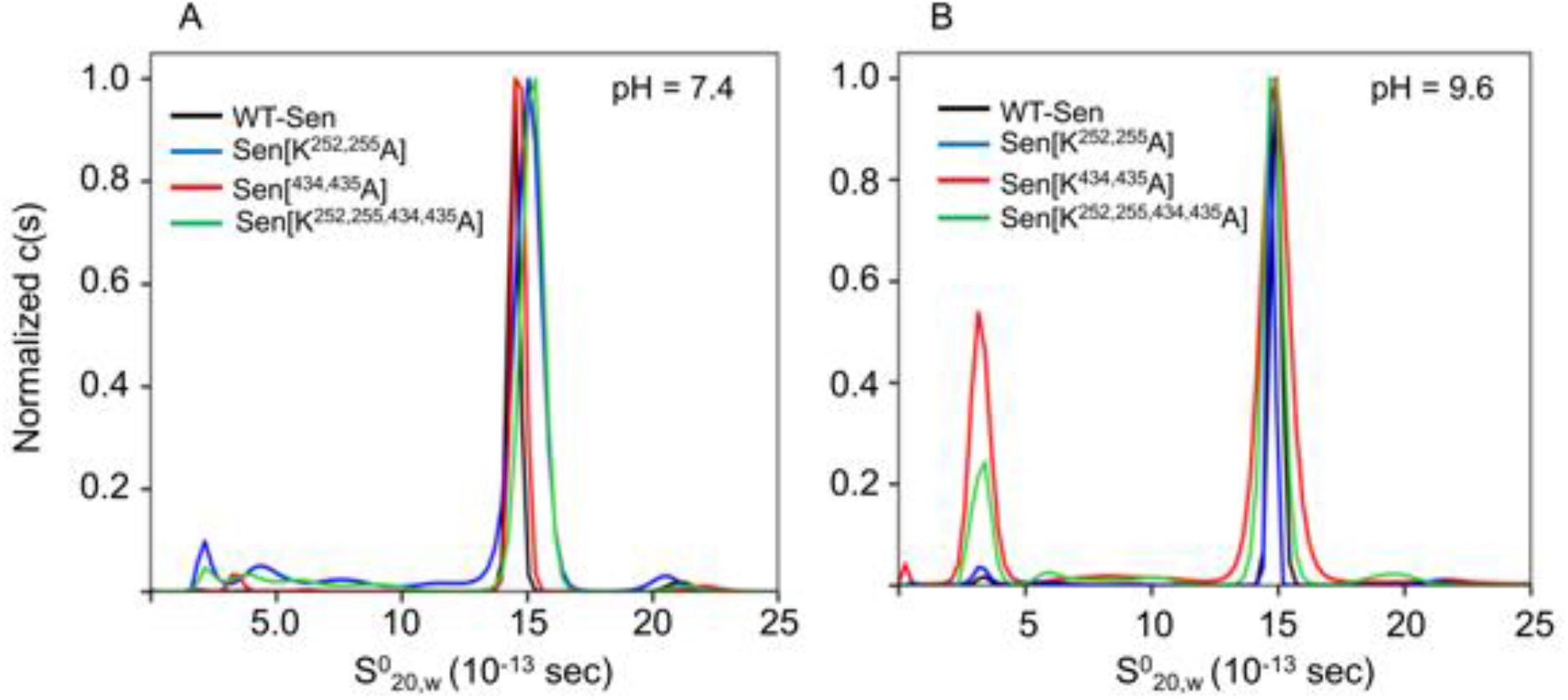
Sedimentation velocities of recombinant SEN mutants. Normalized sedimentation coefficients, c(s), of Sen determined by analytical ultracentrifugation. **(A)** Sedimentation velocity experiments of recombinant Sen mutants were performed at 30,000 rpm, 20° C, in 50 mM Na_2_HPO_4_/100 mM NaCl, pH 7.4. The S°_20,w_ value of each recombinant Sen mutant was overlayed using GUSSI at pH 7.4. Each experiment was performed in triplicate. **(B)** As in **A**, except that the buffer was 50 mM NaHCO_3_, pH 9.6.

### Binding of hPg to Sen Variants

We have shown by sedimentation velocity experiments, as well as isothermal titration calorimetry, surface plasmin resonance (SPR), and ELISA, that hPg does not interact with WT-Sen in solution (20). This is also the case for all of the Sen variants constructed in this study. However, Sen does interact with hPg when Sen adheres to surfaces as shown by similar experiments (9, 20, 28–33). Since previous work had shown that microbial enolases that possess C-terminal lysine residues (27), and also isosteric lysine formed by internal side-chains of K^252^ and K^255^ (34), interacted with hPg through the lysine binding sites of hPg, we investigated whether this was also the case with the highly symmetrical Sen. To establish this point in this work, we investigated by ELISA the binding of hPg to WT-Sen (**Figure 3A**) as well as to the Sen variants that contained Ala mutations at the internal lysine side chains, K^252^ and K^255^ **(Figure 3B)**, at the C-terminal K^434^ and K^435^ residues (**Figure 3C**), and at all four lysines (**Figure 3D**). We find that the C_50_ values for binding of all of the variants to hPg are approximately the same, indicating that hPg binds equally well to the Sen variants that are adherent to the microtiter plate wells. We therefore demonstrate that these previously implicated lysine residues are not directly involved in binding of hPg to octameric Sen in solution or when Sen is immobilized on an artificial surface.

**Figure 3.**
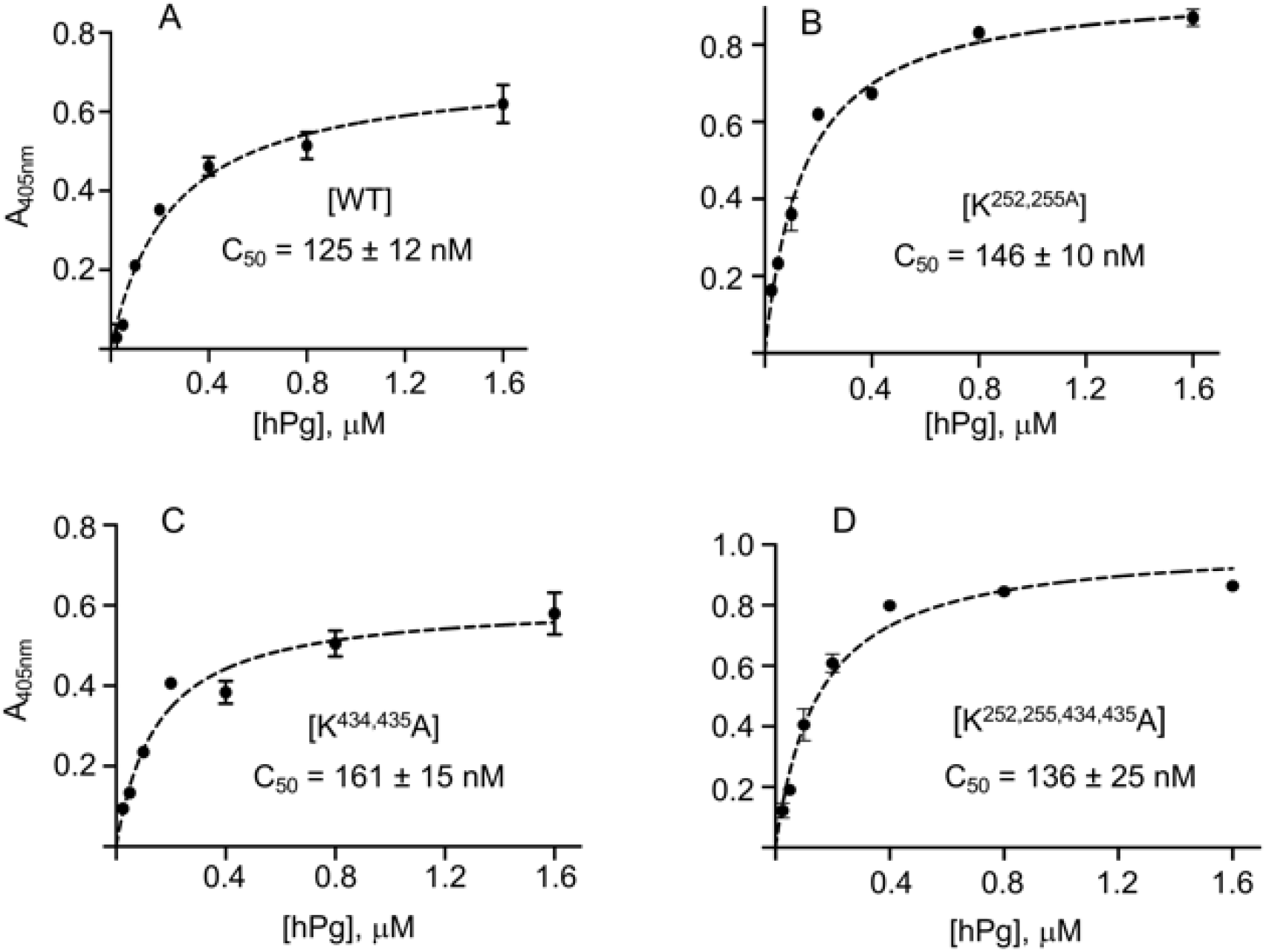
ELISA for the binding of Sen to hPg. Wells of a microtiter plate were coated overnight with the indicated Sen variants (100 μg), after which the wells were washed with PBS and incubated with hPg (0-1.6 μM). After washing with PBS, HRP-conjugated mouse anti-hPg was added and the color was developed with 3,3,5,5’-tetramethylbenzidine. The reaction was stopped by addition of H_2_SO_4_ and the A_450nm_ was determined. The values were plotted against the [hPg] using Prism software and the C_50_ values for each Sen variant were obtained from the fitted plots. The experiments were run in triplicate. When error bars are not visible, they were smaller than the size of the plotted circle.

### Binding of hPg to Sen/DOPG Vesicles

We have previously shown that Sen interacts with DOPG vesicles and that the DOPG-bound Sen enhanced activation of hPg by (tissue-type plasminogen activator) tPA (20). DOPG provides a more biologically relevant system in which to study this interaction with surface-adsorbed Sen. In the present communication, we further demonstrate by flow cytometry with rabbit anti-Sen that the Sen variants constructed herein were equally capable of interacting with DOPG vesicles (**Figure 4A-F**).

**Figure 4.**
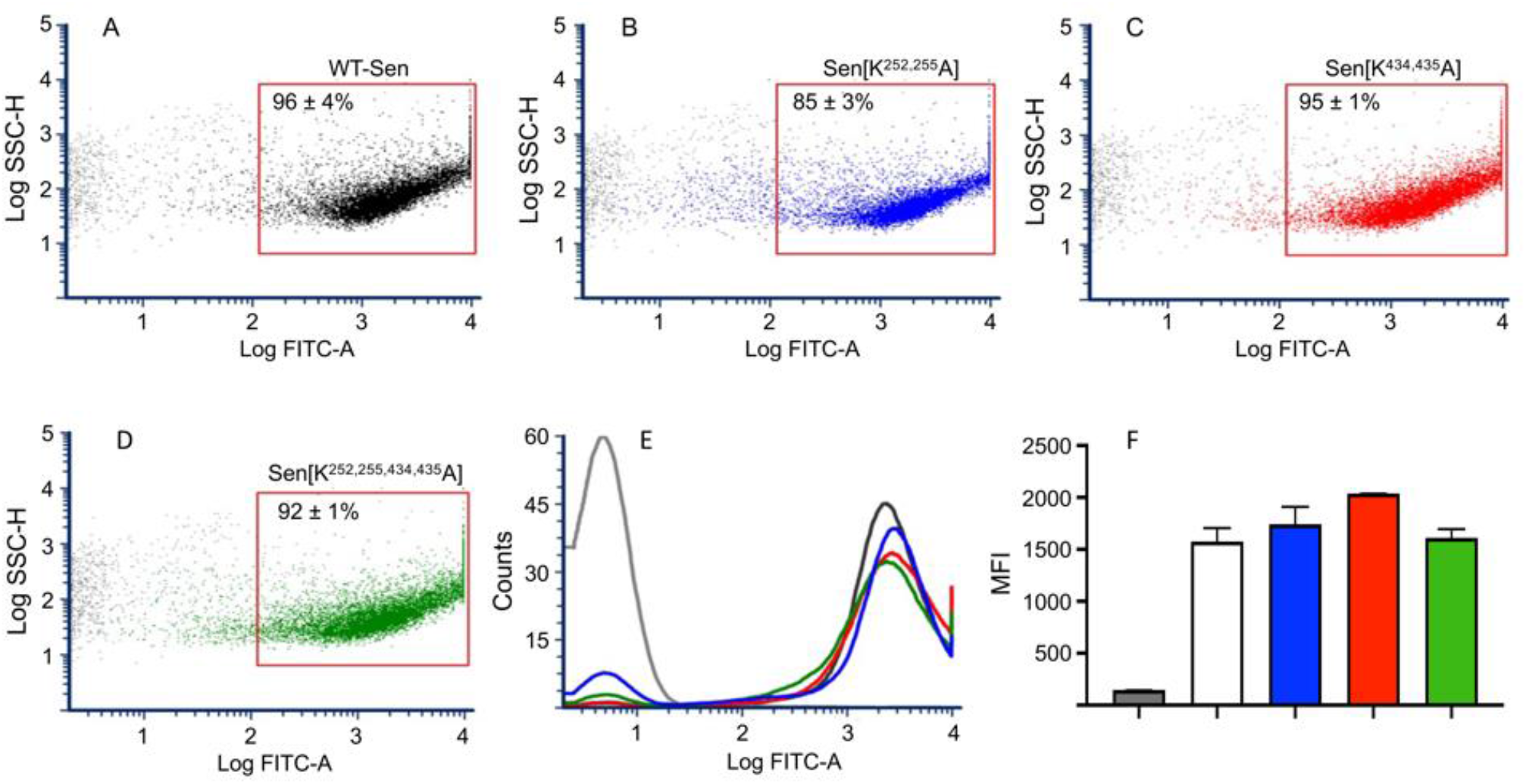
Binding of Sen variants to DOPG vesicles as determined by FCA. The Sen variants were added to DOPG vesicles followed by rabbit anti-Sen and Alexa Fluor 488-chicken-anti-rabbit IgG. **(A-D)** Dot plots showing side scatter (SSC) v*s* FITC fluorescence intensities of DOPG vesicles incubated without WT-Sen (gray) and 5 μM of: **(A)**; WT-Sen (black), **(B)** Sen[K^252,252^A] (blue), **(C)** Sen[K^434,434^A] (red), **(D)** Sen[K^252,255,434,435^A] (green). The FITC-based gate that defines Sen-bound DOPG is indicated on each plot and ranged from 85-95% for all Sen variants. **(E)** Overlay of histograms derived from (**A), (B), (C)**, and **(D)**, showing the distribution of DOPG as a function of FITC fluorescence intensity. **(F)** Bar plots derived from **(E)**, showing the median fluorescence intensity (MFI) of DOPG vesicles.

hPg was added to identical samples of Sen/DOPG as above. Next, anti-hPg was added in place of anti-Sen. Flow cytometric analysis showed that hPg was bound to Sen/DOPG for all Sen mutants (**Figure 5A-F**). These data confirm the ELISA results and demonstrate the binding of hPg to Sen when Sen was adsorbed to different surfaces, including PL. Further, all of the DOPG/Sen variant/hPg complexes were equally activated by (tPA) (**Figure 5G,H**). These data show that not only were the Sen variants fully capable of binding to PL vesicles but that hPg on these DOPG/Sen variant complexes was functional and fully activatable by tPA. This confirms that mutation of these particular lysine residues of Sen did not attenuate hPg binding or activation on a PL vesicle.

**Figure 5.**
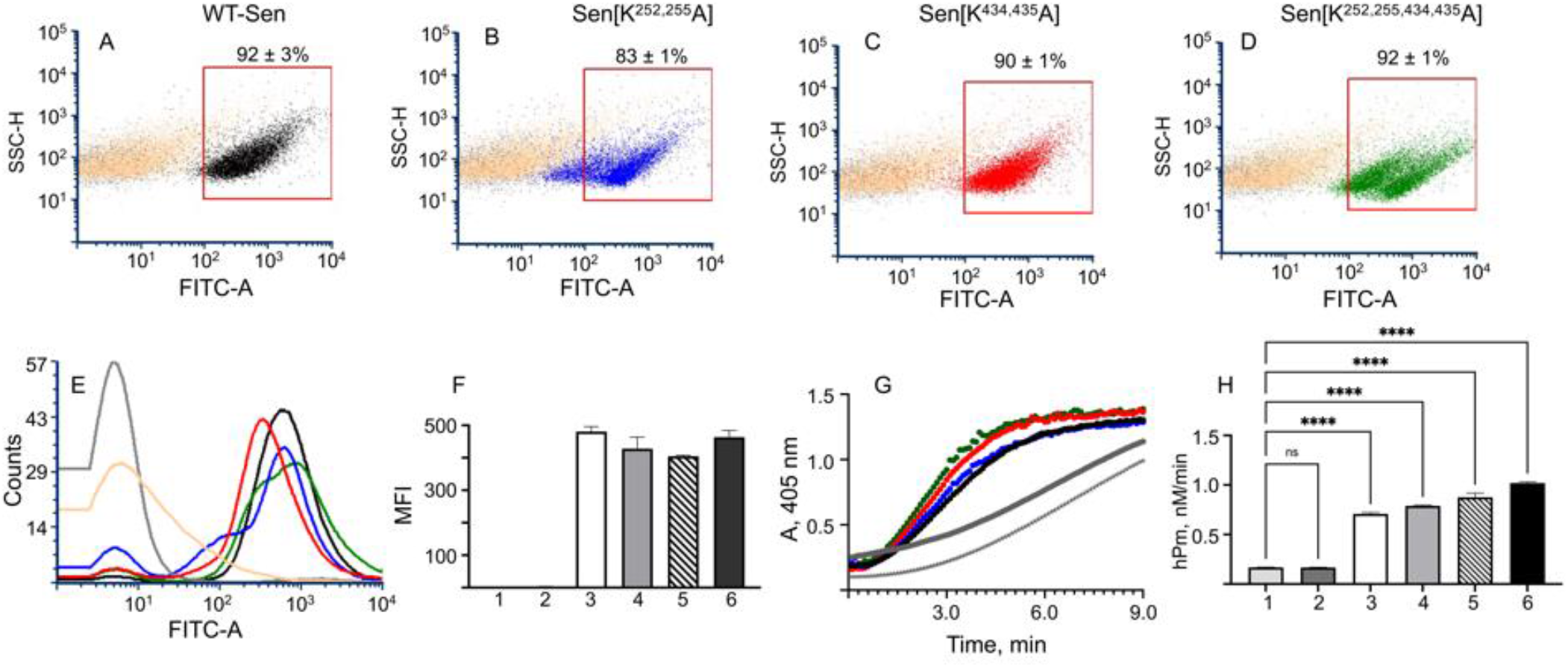
Functional acquisition of hPg by Sen/DOPG vesicles. **(A-D)** Dot plots showing side-scatter (SSC) vs FITC fluorescence intensity of: **(A)** DOPG/WT-Sen/hPg (black); DOPG/Sen[K^252,252^A]/hPg (blue); **(C)** DOPG/Sen[K^434,434^A]/hPg (red); and **(D)** DOPG/Sen[K^252,255,434,435^A]/hPg (green) overlaid on those of DOPG/no Sen/hPg (gray) and DOPG/WT-Sen/no hPg (tan) as determined by FCA. A final concentration of 0.8 μM hPg was added to replicate DOPG/Sen complexes prepared as in **Figure 4**, followed by mouse anti-hPg, and Alexa Fluor 488-donkey-anti-mouse IgG. The FITC-based gate that defines DOPG/Sen bound to hPg is indicated on each plot and ranges from 83-93%. **(E)** Overlay of histograms derived from **A, B, C**, and **D**. **(F)** Bar plots of median fluorescence intensities (MFI) derived from **E**: (1) DOPG/no Sen/hPg; (2) DOPG/WT-Sen/no hPg; (3) DOPG/WT-Sen/hPg; (4) DOPG/Sen[^K252,255^A]/hPg; (5) DOPG/Sen[K^434,435A^]/hPg; and (6) DOPG/Sen[K^252,255,434,435^A]/hPg. **(G)** The rates of hPg (0.2 μM) activation by tPA (3.5 nM) in the presence of DOPG/no Sen (gray circle); no DOPG/no Sen (gray asterisk); DOPG/WT-Sen (black); DOPG/Sen[K^252,252^A] (blue); DOPG/Sen[K^434,434^A] (red); and DOPG/Sen[K^252,255,434,435^A] (green). **(H)** The initial velocities of hPg activation calculated from the linear regions of plots of A405 *vs* t^2^. (1) no DOPG/no Sen/hPg; (2) DOPG/no Sen/hPg; (3) DOPG/WT-Sen/hPg; (4) DOPG/Sen[K^252,255A^]/hPg; (5) DOPG/Sen[K^434,434^A]/hPg; and (6) DOPG/Sen[K^252,255,434,435^A]/hPg. Pairwise comparison performed between (1) and (2 to 6) provided probability (p) values as indicated by asterisks. **** p< 0.0001, and ns, not significant.

### Cryo-EM-based High-resolution Structure of WT-Sen

The micrographs of Sen that were collected in the cryo-EM analysis depicted only an octameric structure present in a variety of different orientations (**Figure 6A**). A small percentage of stacked dimers of octamers was observed, but these structures constituted <1% of particles and were therefore omitted from data processing. A total of 3,258 micrographs were curated manually and 2,756 micrographs were chosen for particle selection, which was performed using blob selection with a minimum radius of 50 Å, maximum radius of 150 Å, and box size of 256 pixels at a pixel size of 1.29 Å. This resulted in ~2,000,000 particles. Manual selection was performed to input Contrast Transfer Function (CTF) parameters and remove potentially incorrectly selected particles. A 2D classification of the initial particle set was performed using 40 iterations with a maximum resolution of 6 Å. Of the 50 classes obtained, 10 were selected and used for template particle selection. This resulted in a refined particle set consisting of 813,556 particles. Fifteen classes were chosen, the 2D classification was again performed, and 17 final classes (**Figure 6B**) were selected consisting of 442,256 particles in the final count. Each orientation was stacked with particles that differed by no more than 5°. *Ab initio* reconstruction was performed using the final particle set to obtain a 3D map structure which was then refined using homogeneous refinement. This created an initial 3.1 Å density map of octameric enolase. This map was then refined by applying an experimentally-generated 3D mask with point group 422 (D4) symmetry (EMDB 26406), as shown in **Figure 6C**. A visual representation of the workflow provided here are presented in **Supplemental Figure 1S**.

**Figure 6.**
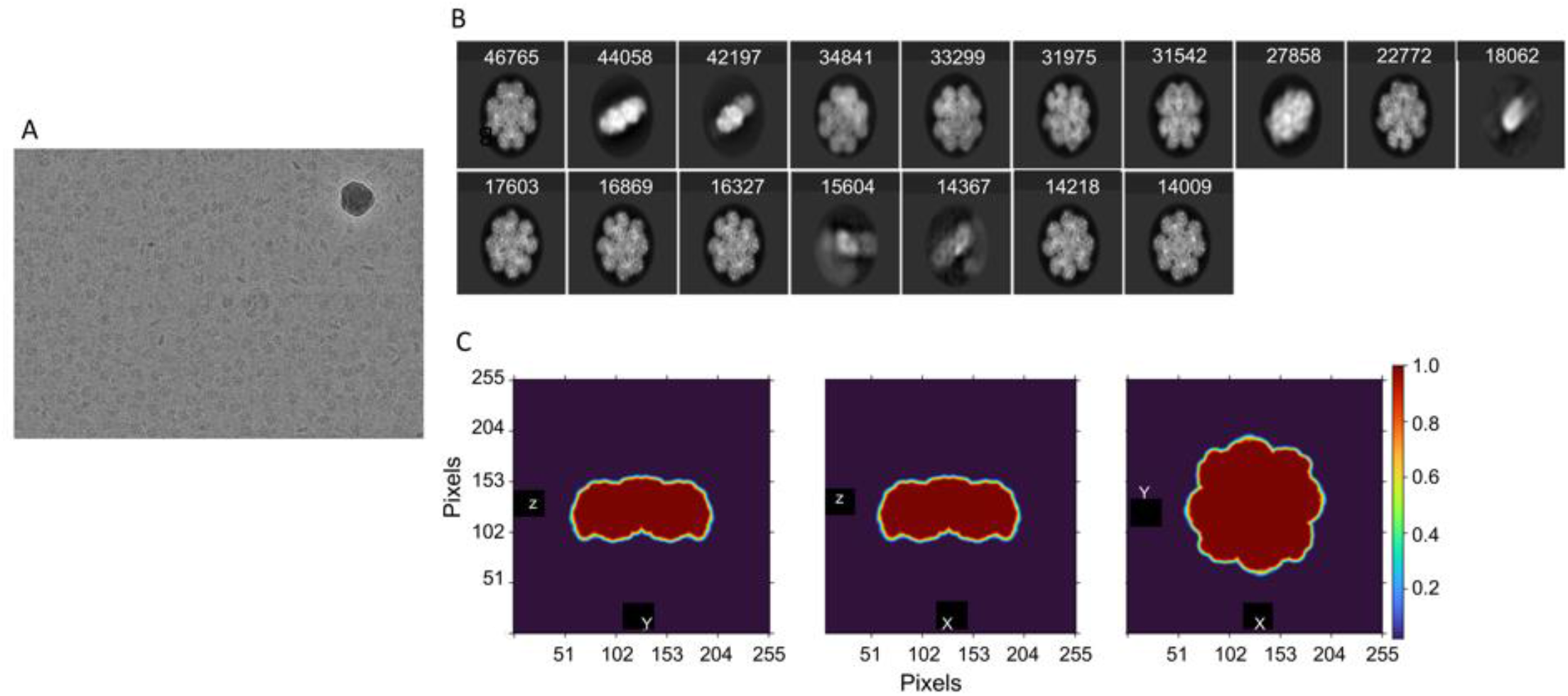
Progression of cryo-EM map construction. A scheme showing the process of generation of the cryo-EM map from the images taken using the Titan Krios. **(A)** Micrographs of Sen that shows the octamer in different orientations. The crystalline ice in the top right corner is not considered. **(B)** Particle images used to obtain final 2D classes, taken as an average of 442,256 particles. The number of patricles (± 5°) in each orientation is provided above the 17 final images. Autosharpened density maps of the 3D structure of Sen refined from **(B)** to 2.6 Å that show the overall structure of the surface of the octamer. Panels 1) and 2) are maps rotated 90° around the z-axis showing the high internal symmetry of Sen, thus allowing imposition of D4 symmetry.

The final map was sharpened using Phenix 1.20 AutoSharpen map function to obtain a final resolution of Sen at 2.6 Å. All resolutions provided for the cryo-EM map were measured using Fourier-Shell Correlation (FSC) at the highest standard FSC 0.143. FSC is a measure of the correlation between our finalized map and corresponding half maps to prevent over-correction or structural artefacts by the applied mask. The FSC plot for the finalized map is provided in **Supplemental Figure 2S-A**.

### Molecular Modeling of Sen

The 2.6Å resolution map was used as the template for construction of the model for Sen (PDB 7UGU). The map was divided into monomeric sections for initial modeling in the Phenix 1.20 Map-to-Model algorithm. This model was imported into ChimeraX and complete sections of the model were placed in space using the experimentally determined map structure as a template along with replacement of residues using the amino acid sequence of Sen from GAS isolate AP53 (PDB 3ZLH). Once the amino acids were corrected, the structure was real space refined in Phenix Fit-to-Map. This process was repeated until the fully modeled monomeric structure was constructed. After the model of the monomer was completed, it was input into Phenix to dock into the map using the known octameric symmetry file generated in Phenix. Since it is known that Sen in solution is an octamer, we constructed a solution model of Sen in order to more fully represent how Sen assembles as an octamer in solution. This structure was real space refined to align it in the highest fit within the density of the cryo-EM map. This fit was then measured using Map *vs* Model FSC and determined at a final resolution of 2.8 Å **(Supplemental Figure 2S-B)**. The secondary structure of the octamer was confirmed using a Ramachandran plot with PDB validation. After using Phenix and ISOLDE in ChimeraX (35), the final model shows 94.9% favored, 5% allowed, 0.1% outliers and an overall clash score at 0.02 (**Supplemental Table 2S**).

### Structurally Significant Residues in the Major and Minor Interfaces of the Octamer Interfaces

The final 2.6Å structure of the Sen homooctamer is shown in cryo-EM map **(Figure 7A,B)** and ribbon structures with imposed electron densities (**Figure 7C,D)**. The monomers are shown in different colors to represent each protomeric subunit, A-H. The model shows that the Sen octamer consists of eight monomers which assemble to create four major interfaces (A-B, C-D, E-F, and G-H) and four minor interfaces (B-C, D-E, F-G) between the monomers, with the circular structure completed by the minor interface at monomers H-A. The buried surface areas in the major and minor interfaces were 3560Å^2^ and 2470Å^2^, respectively (https://www.ebi.ac.uk/pdbe/pisa). Each monomer contains two definable regions. The NH_2_-terminal domain consists of three antiparallel β-strands, followed by an unstructured loop (L) region comprising residues V^37^/K^56^ (L1 in **Figure 8A,B**) and four α-helical regions. The C-terminal domain primarily consists of the Sen active site formed by a highly stable 8-stranded β/α barrel surrounded by a series of seven α-helices, similar to that found in the X-ray structure (36). Two additional relatively unstructured loops are observed, comprising residues G^152^/Q^163^ (L2 in **Figure 8A,C**), and Y^250^/T^260^ (L3 in **Figure 8 A,D**). Close examination of the major interfaces formed between two antiparallel subunits and the minor interfaces allowed us to highlight the residues that are likely involved in stabilization between Sen subunits in the major interface (**Figure 9**) and the Sen subunits in the minor interface (**Figure 10**) in solution. The residues at the major interfaces with appropriate electrostatic interactions are R^58^/E^177^, R^401^/D^403^, and R^10^/D^416^ (**Figure 9B)**. Close-up views of the major interfacial hydrogen bonds between S^400^/R^401^ and S^15^/T^402^/D^403^ (**Figure 9C**), and Y^8^/E^22^/R^413^/D^416^ (**Figure 9D**) are shown.

**Figure 7.**
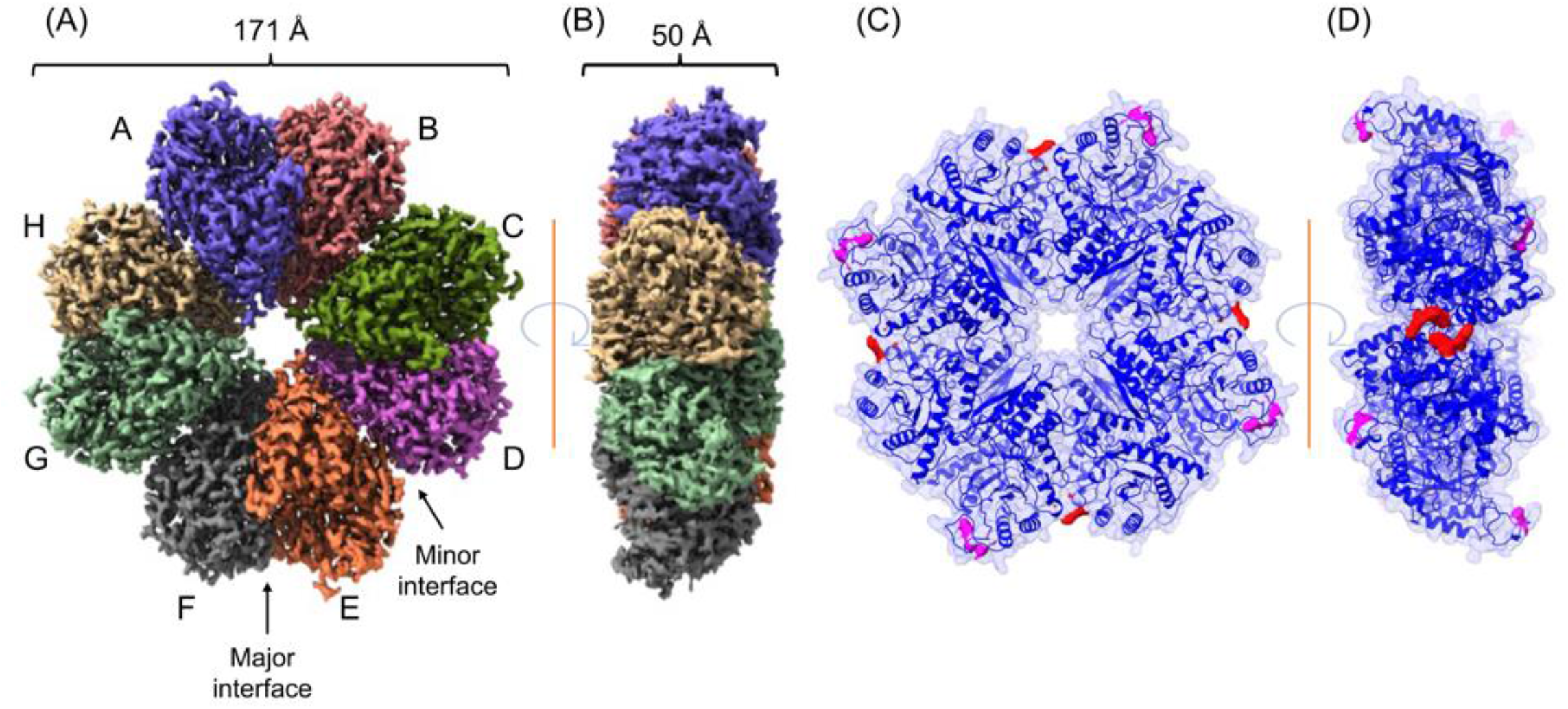
The octameric structure of Sen as determined by the dimensions of the experimental cryo-EM map. The cryo-EM map reconstruction of sharpened octameric Sen is divided into the eight identical subunits (A-H) of monomeric Sen from the **(A)** front (171 Å) and **(B)** side (50 Å) views. The structure consists of major interfaces (monomers A/B, C/D, E/F, G/H), and minor interfaces (monomers B/C, D/E, F/G, H/A). **(C, D)** The ribbon structure of Sen highlighting the secondary structure of the complex overlayed with the model surface in **(C)** front and **(D)** side views. The location of Loop-3 (L3) in one of the monomers is indicated. Also shown are the side-chains previously proposed to be involved in the binding of Sen to hPg. However, K^434^ and K^435^ (red) are buried in the octamer facing inward to stabilize the major interfaces of the octameric structure. K^252^ and K^255^ (pink) are exposed on the surface on the front-facing L3 of Sen.

**Figure 8.**
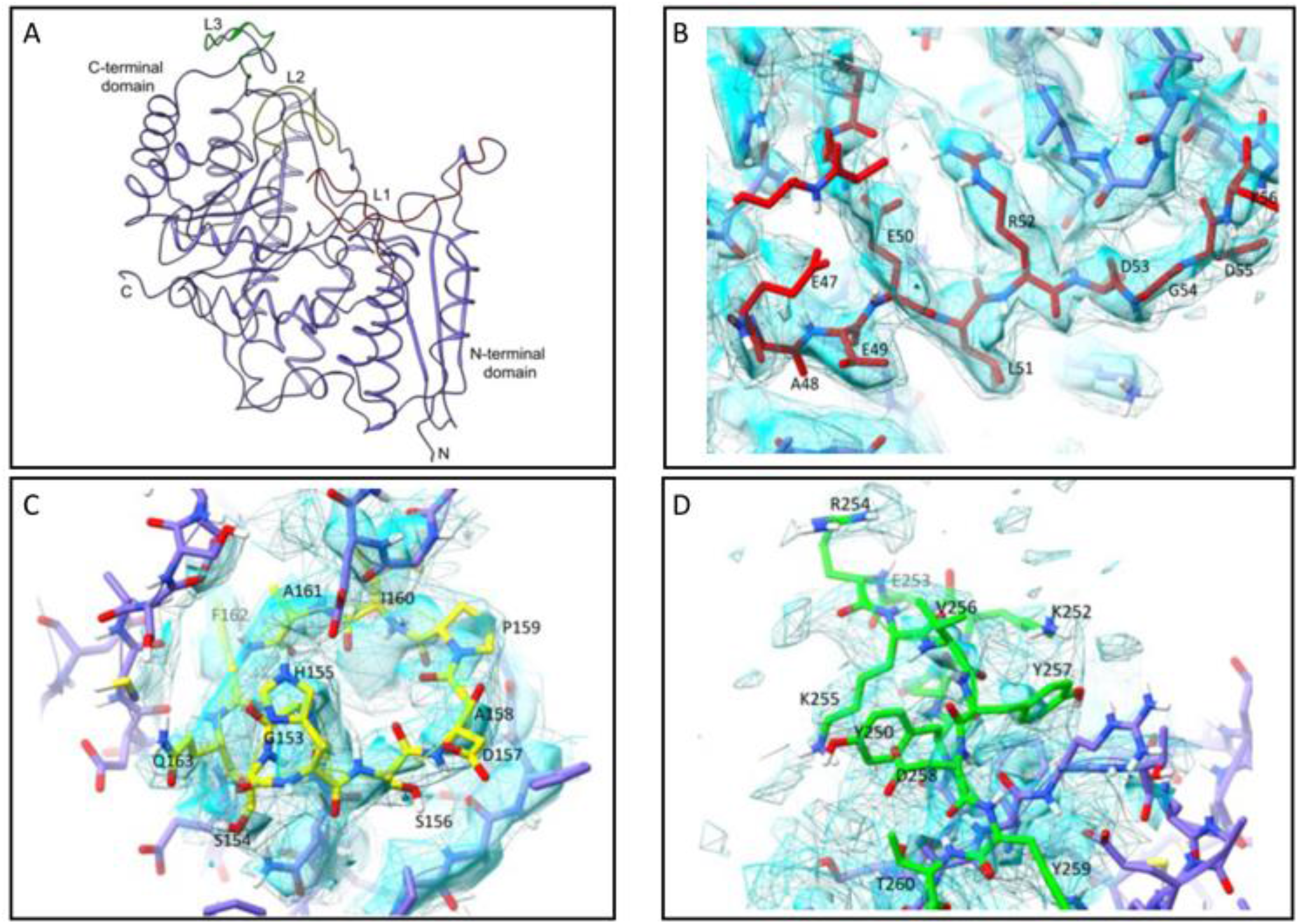
3D structure of Sen. **(A)** The monomer subunit of the octameric Sen exposing loop 1 (L1; red), loop 2 (L2; yellow), and loop 3 (L3; green). Close-up view of the cryo-EM map in the loop regions L1 (B), L2 (C), and L3 (D). L3 has the poorest electron density and the most flexible structure compared to the L1 and L2.

**Figure 9.**
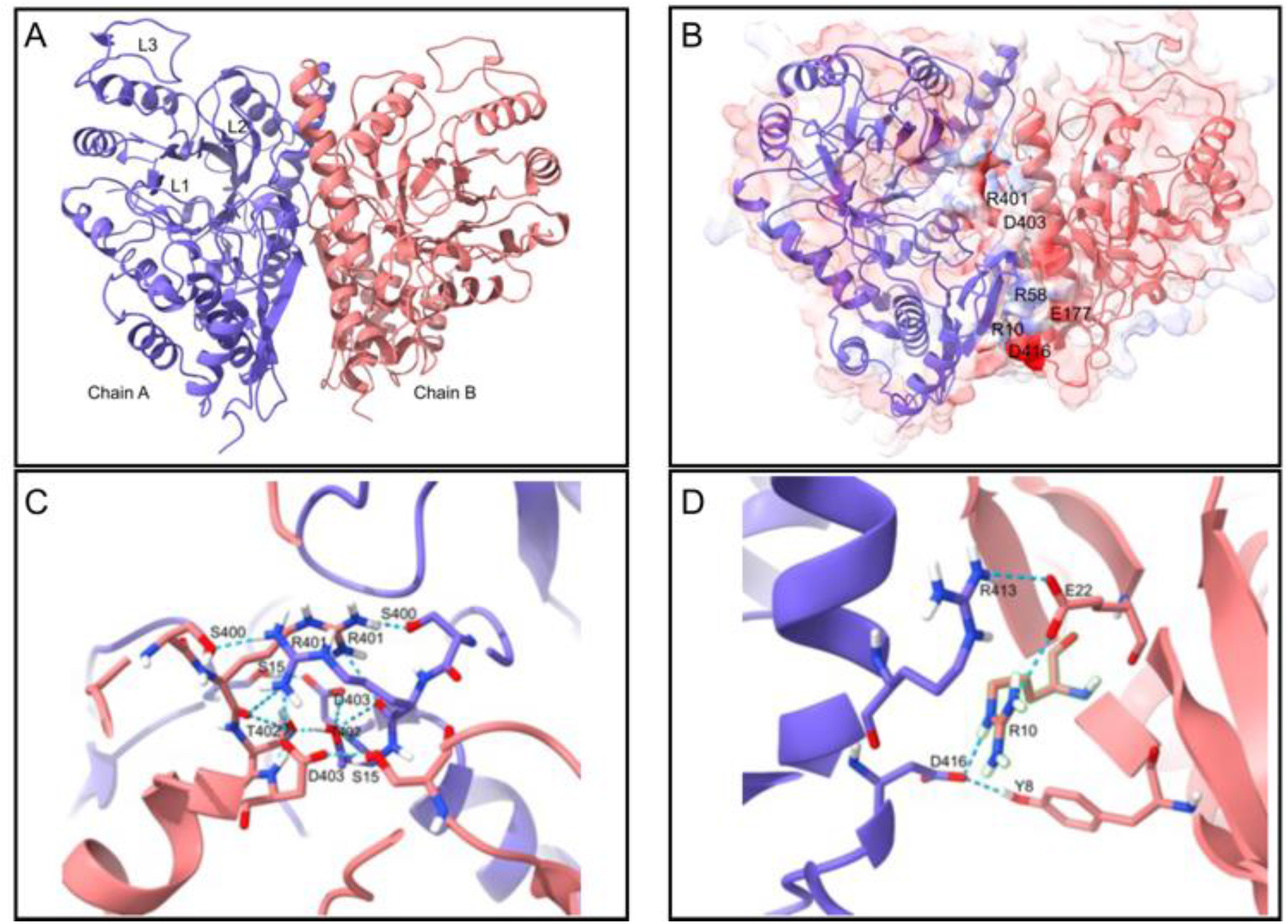
Amino acids primarily responsible for stability at the major interface. The major interface associates by hydrogen bonded and ionic residues with distances between 2.6-3.0Å. **(A)** The overall ribbon structure between the major interfaces. The positions of the three loops (L) are indicated. **(B)** Electrostatic interactions that help stabilize the major interface, specifically, R^58^/E^177^; R^401^/D^403^; and R^10^/D^416^. **(C-E)** The close-up view of hydrogen bonds between residues; **(C)** S^400^-OG/R^401^-NH1; S^15^-OG/D^403^-OD^1^/T^402^OG; **(D)** R^413^-NH1/E^22^-OE1; D^416^-OD2/Y^8^-OH; D^416^-OD1/^R10^-N; and R^10^-NH1/E^22^-OE1.

**Figure 10.**
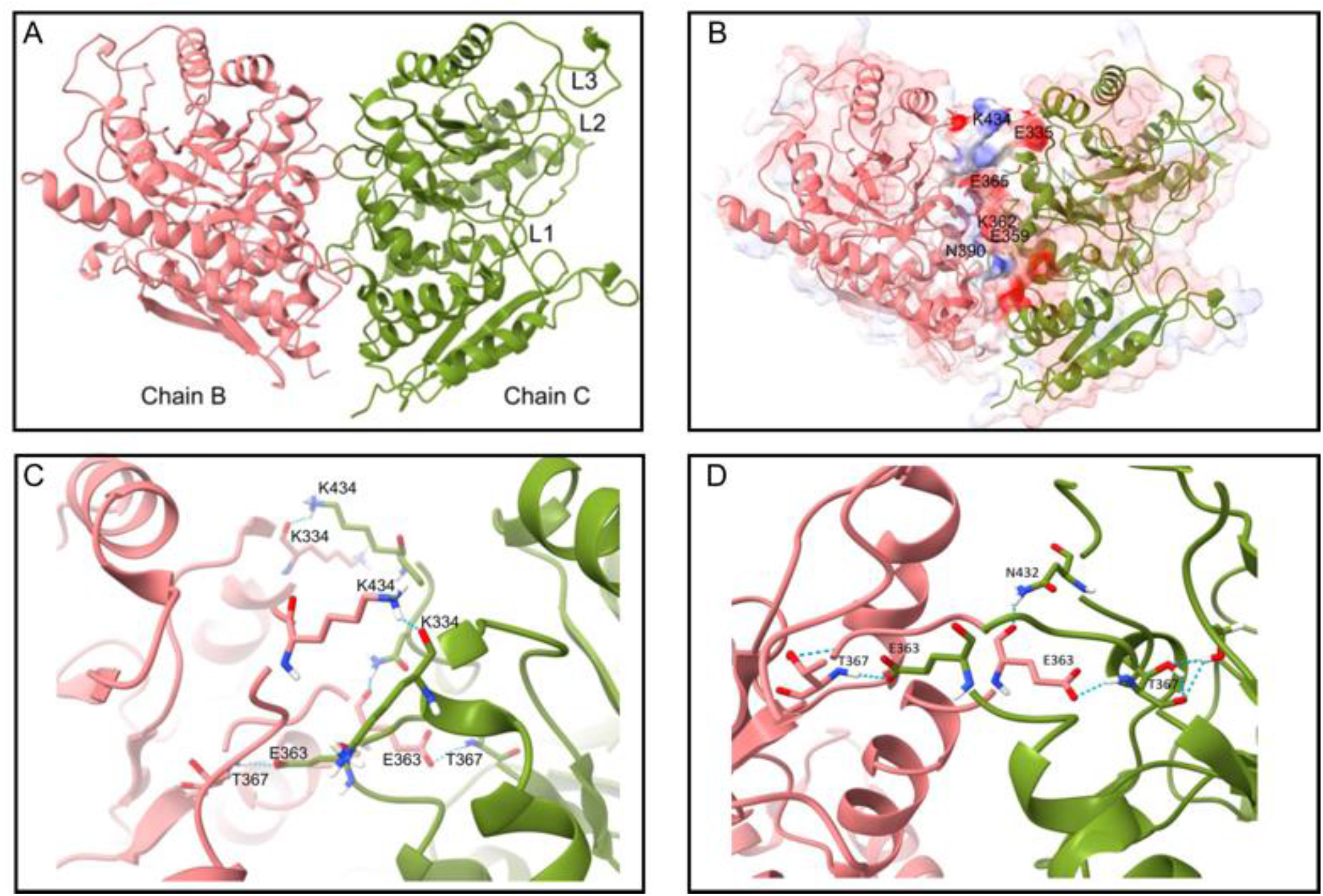
Amino acids primarily responsible for stability at the minor interface. The minor interfaces of the octamer associate by hydrogen bonded and ionic residues with distances between 2.6-3.0Å. (A) The overall structure ribbon structure between the minor interfaces. The locations of the three loops (L) are indicated. (B) Partially-exposed electrostatic contact surfaces between K^434^-^/^E^335^; K^362^/E^365^; and E^359^/N^390^ that help stabilize the minor interface. **(C)** Close-up view of the hydrogen bonds between residues K^434^-ND2/K^334^-O and T^367^-N/E^363^-OD2. (D) Close-up view of the hydrogen bonds between residues N432^ND2^/E363^N^; and E363^OE1,OE2^/T367^N^.

A ribbon structure of one of the minor interfaces is shown in **Figure 10A**. The residues likely involved in the electrostatic stabilization of these interfaces are K^434^/E^335^;K^362^-E^365^, and E^359^-N^390^. (**Figure 10B**). Close-up views of the interfacial hydrogen bonds between K^334^/K^434^; and E^363^/E^367^ (**Figure 10C**), as well as E^363^/T^367^ and E^363^/N^432^ (**Figure 10D**), are shown. Importantly, the COOH-terminal residues, K^434^ and K^435^ exist within the minor interface and not on the surface as shown by the hydrogen bond of the *ε*-amino group of K^434^ and the amide of K^334^ between these residues (**Figure 10C**).

### Comparison of the Cryo-EM and X-ray Crystallographic Structures of Sen

While the technology used to obtain the high-resolution Sen structure by these two methods is highly disparate, some comparisons can be made. Both structures exhibit key similarities in the thickness of octameric structure as well as each subunit structure, such as the helixes and strands. Key differences between the previously reported X-ray crystal octamer and the current single particle cryo-EM structure lies in the total size. The diameter of the Sen octamer is 171 Å, which is larger than the diameter of 153 Å previously modeled from X-ray crystal structure data (36). The difference of approximately 800 Å^2^ buried surface between the crystal structure and the cryo-EM structure is most likely due to the nature of the crystal packing constraints and solvent composition of the Sen X-ray crystal structure (36). The increase in diameter is primarily due to the extension of the L3 loop into solution which occurs by way of interactions with the surrounding water molecules in the cryo-EM structure, as opposed to being tucked-in in the crystal structure. This extension into solution allows L3 to fulfill its proposed role of participating in enolase activity by closing around the substrate (36). Moreover, the cryo-EM structure has ~100 Å^2^ less buried surface area, at each of the major and the minor subunit interfaces

### Sen Previously Proposed hPg Binding Sites

Past studies have proposed that the Sen C-terminal residues (K^434,435^) was believed to be hPg binding sites (9, 18, 37). A more recent study also suggested that the internal lysine residues in L3 (K^252,255^) play a role in hPg binding in a concerted manner with the COOH-terminal lysine residues (18, 31, 34, 36), an observation that was also suggested using molecular docking between hPg and the Sen minor interface (36).

In comparing the structure of our model of Sen in solution and the crystal structure of monomeric Sen, one of the prominent differences is observed in the important loop 250-260 of Sen. The loop is seen to jut out in solution due to its hydrophilic nature (**Figure 7C,D**), a factor that further stabilizes the octameric structure with the surrounding water molecules or the charged cell wall surface. This loop was also suggested to play a role in stabilizing the enolase enzyme activity by closing the active site once it interacts with substrate (36). If this is the case, this infers that Lys^252,255^ may not play a role in hPg binding in its native state in solution. Additionally, the COOH-terminal lysine residues (**Figure 7C,D)** are located between the minor interface making them inaccessible for hPg binding. In accord with these observations, we note from our direct binding measurements that neither K^252^, K^255^, K^434^, nor K^435^ directly participate in the binding of hPg to Sen in solution, or when Sen is bound to surfaces. There are many additional surface lysines in Sen with four showing the characteristic Lys, Glu helical i, i±3 **(Figure 11A-C)** orientation shown previously to be a binding site for hPg. These are potential targets to be explored for their viability and ability to bind hPg on the cell surface.

**Figure 11.**
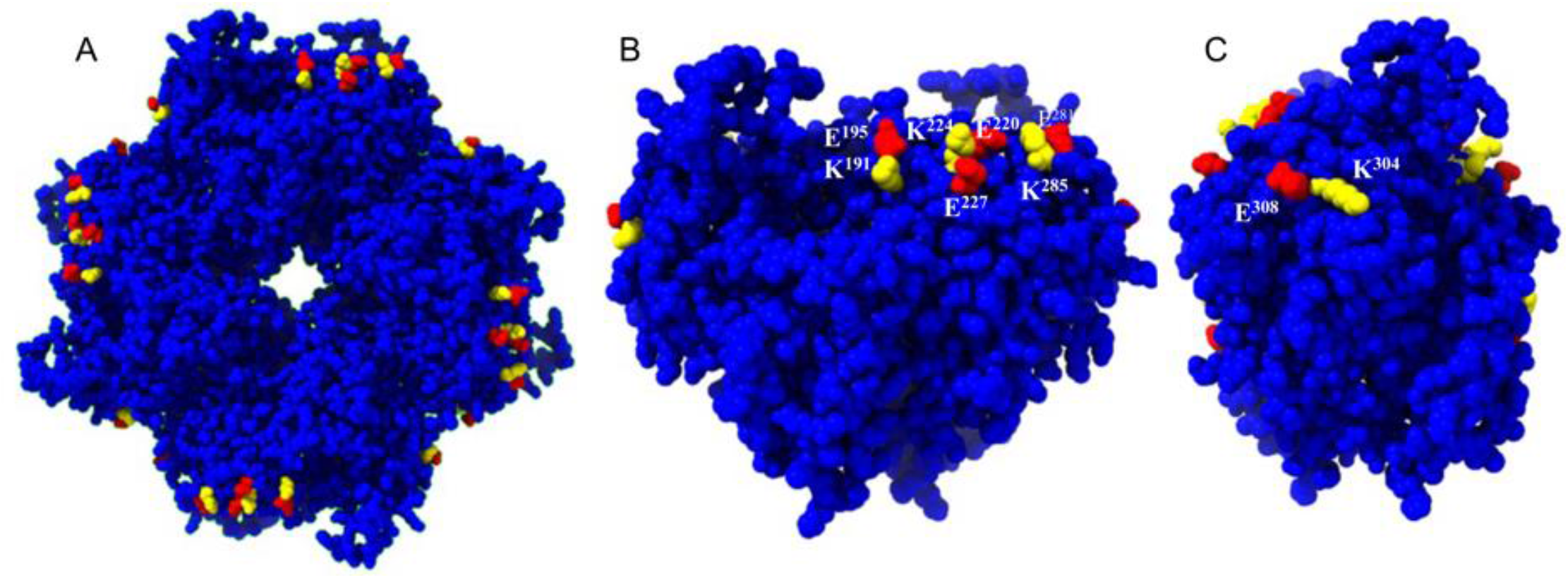
Surface lysine residues with potential to bind hPg. (A) The surface of the Sen octamer with helical Lys side-chain residues in yellow and i, i±3 Glu residues in red, located on the surface of the protein. These Lys residues aid in binding of the Sen to the negatively charge sections of the cell wall and lysines with an i, i±3 glutamic acid in a helical region of the protein have been previously characterized as potential binding sites for hPg. (B) The surface of a single monomer of the homooctamer. Lys residues are labelled yellow and Glu residues are labeled in red (their sequence numbers are shown). (C) Lys and i, i±3 Glu residues in the minor interface are postulated to be potential binding sites for hPg as well as providing stability to the octamer as a whole.

### Octameric Instability of [K^252,255,434,435^A] Mutant Leads to Hexamer, Heptamer, and Tetramer Formation

As we have proposed, the function of the K^434^K^435^ residues is primarily in stabilizing the octameric structure of Sen. Generation of a cryo-EM map (EMDB:27407) and model (PDB:8DG4) for the [K^252,255,434,435^A] mutant showed that the removal of the C-terminal lysines led to the octamer becoming much more unstable. The 2D classifications of the sample showed the existence of incomplete Sen octamers in the form of tetramers, hexamers, and heptamers **(Figure 12)** that were not present in WT-Sen. These structures make up 15% of the total particles imaged and were likely a result of the octameric instability due to the diminished level of interactions in the minor interface. Hydrogen bonding interactions with K^334^ and electrostatic interactions with E^335^ with these C-terminal lysines stabilize the octamer in WT-Sen **(Figure 9B)** and breaking this interaction leads to increased flexibility at the minor interface in Sen[K^252,255,434,435^A], plausibly resulting in a single/multiple monomeric subunits dissociating from the octamer and making the remaining octameric Sen[K^252,255,434,435^A] more flexible. This higher flexibility leads to the lower resolution (3.1Å) of the Sen[K^252,255,434,435^A] map due to higher variability in the particles themselves (PDB:8DG4).

**Figure 12.**
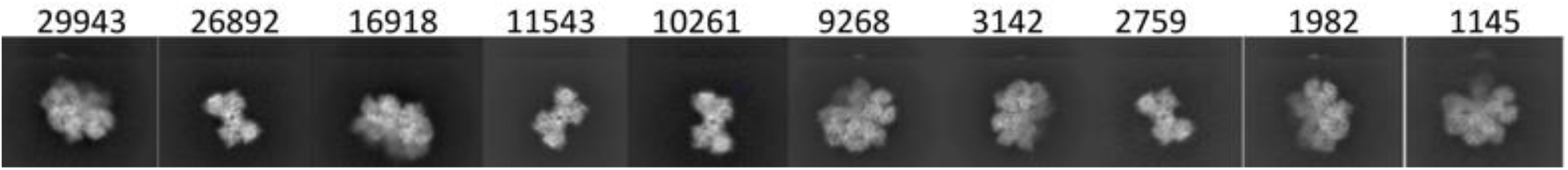
Two-dimensional (2D) classification of Sen[K^252,255,434,435^A] instability. Mutation of the C-terminal lysines results in octameric instability that decreases the percentage of octamer to 85%. The remaining 15% consists of a combination of monomers, tetramers, hexamers, and heptamers, with the numbers of each image ± 5° shown above each image. The mutant also demonstrates an increase in octamer flexibility that could allow for increased exposure of amino acids within the minor interface. The above classifications make up 15% of the total ~800,000 particle dataset collected.

## DISCUSSION

hPg receptors/binding proteins (hPg-R) are present on most eukaryotic and prokaryotic cells and serve to bind hPg, stimulate its activation to hPm, and to protect the cell-bound hPm from inactivation by natural inhibitors, *e.g*., α2-antiplasmin. This formed and stabilized potent serine protease on cells assist in their migratory functions that are needed for processes such as macrophage recruitment during inflammation (38, 39), fibrin surveillence (40), wound healing (41), neurite outgrowth (42), and cancer cell growth, invasion, and metastasis (43–45), among others. In the case of prokaryotes, *e.g*., GAS, the hPm generated on the cell surface allows microorganisms to evade immune responses of the host in various manners, *e.g*., resistance to complement-mediated opsonization (46, 47) and resistance to killing by extracellular histones (48). In sum, microbial-bound hPm transforms the cells into proteolytic microorganisms by conscripting and activating host hPg, thus allowing the microbes to digest cellular protein barriers, *e.g*., the extracellular matrix, that inhibit their dissemination and metastasis.

The focus of this communication is the nature of the surface receptors on GAS cells. At least three such receptors/binding proteins have been identified, specifically the endogenous GAS M-protein, and the moonlighting glycolytic enzymes Sen (9) and glyceraldehyde-3-phosphate dehydrogenase (GAPDH) (49). The M-protein of the highly prevalent Pattern D GAS strains (PAM) is the only M-protein that binds hPg and hPm directly and tightly (50). In these strains, a subform of the endogenous hPg activator SK, *viz*., SK2b, that activates hPg maximally when hPg is bound to PAM, is coinherited with the PAM type M-protein (51). As a further consideration, some Pattern A M-proteins contain a strong binding site for fibrinogen. This M-protein/fibrinogen complex can also bind hPg and is converted to hPm by another more general coinherited subclass of SK (SK1 or SK2a), which activates hPg in the M-protein/fibrinogen/hPg complex and also activates hPg in solution (14, 15, 52).

Bacterial enolase-type Pg-Rs, *e.g*., Sen, contain the requirements to serve in this capacity. First, Sen is exposed on the cell surface. This protein is transported from the cytosol, where it serves its glycolytic function, to the cell surface without benefit of a signal polypeptide or other known targeting sequences. After transport, Sen is bound to the outer cell membrane of Gram-negative bacteria. In Gram-positive bacteria, Sen is bound to the cell wall and exposed to the surface (20). Secondly, many, but not all, enolases possess a C-terminal lysine residue, a feature believed to be critical to this protein serving the function of a Pg-R, since in this manner enolase can bind to the lysine binding sites of hPg in an EACA dependent manner.

We tested these concepts in this study. Sen in solution does not interact with solution phase hPg; however, surface-bound Sen, whether adsorbed to SPR chips, microtiter plates, GAS cell surfaces, or phospholipid vesicular surfaces, interacted with hPg with a Kd of ~100-200 nM (20). The binding of hPg to Sen is weaker than that of hPg to the primary GAS Pattern D PgR, PAM, which is ~1 nM, but in strains without PAM it is possible Sen is an important tool for recruiting hPg to the surface. Using PL (DOPG) vesicles in which Sen was embedded, leads us to believe that we have an important membrane model for studying various aspects of Sen-hPg binding.

Several studies have been published which demonstrated that alteration of the two C-terminal lysine residues, K^434,435^, eliminated hPg binding to Sen adsorbed to cell surfaces (27, 34). We did not observe this behavior and show herein that hPg interacted with adsorbed Sen[K^434,435^A]/PL with equal affinity as WT-Sen/PL. Further, the tPA-catalyzed activation of hPg adsorbed to Sen[K^434,435^A]/PL occurred to approximately the same extent as hPg adsorbed to WT-Sen/PL. The same was found to be true for hPg binding to Sen[K^252,255^A], another set of Lys residues believed to be important in the interaction of hPg and Sen (34), and to the mutant with all four Lys residues modified, *viz*., Sen[K^252,255,434,435^A]/PL. It is stressed that Eno is assembled in various quaternary structures depending on the species from which the protein has been obtained. In mammalian organisms and in yeast, enolase is dimeric (53), whereas *E. coli* enolase is a hexamer (54, 55). In streptococcal and staphylococcal species, enolase is a very symmetrical doughnut-shaped homooctamer, composed of four homodimers (31).

Crystal structures of enolase from several species are provided in the PDB database. These structures in general suffer from some generic well-known limitations, such as crystal dependency of the structure, crystal lattice packing constraints, interactions with components of the mother liquor, and the fact that crystal structures provide a static snapshot of the structure in the crystal under very specific crystallization conditions. Using cryo-EM, we created an experimental map of the full octamer of Sen in solution. We then modelled a monomer from a section of the map and fit the monomer to form an octamer in the original experimental map, so that it accurately reflected the full octameric structure in the dimensions determined by the experimentally-determined map. This allowed us to model the interactions of octameric enolase in solution.

Creating a model based on cryo-EM analysis provides an opportunity to better consider the structure of Sen in solution in ways that are not represented in crystal structures. These X-ray structures also do not demonstrate how molecules orient in solution in the formation of the octamer complex. The benefit of our cryo-EM model is that it directly demonstrates the structure of enolase as it exists in solution and is built from 15 different octameric conformations comprised of >400,000 unique particles that are frozen during vitrification, thus providing elements of a dynamic structure. We built sections of the model directly from the cryo-EM map and then assembled these sections within the map based on how they are oriented in the enolase octamer, as opposed to having crystallized monomers or dimers. Thus, we provide insights into the manner in which the subunits of Sen assemble in solution. We were able to more clearly display residues R^254^-K^255^, within L3, and the COOH-terminal lysine side chains, K^434^-K^435^, that were not visible in the Sen crystal structure. Since these regions have been proposed to interact with hPg, the ability to clearly display them helps to further advance studies in the field.

The most frequently cited areas of Sen for its interaction of hPg is centered on the K^434^K^435^ C-terminal residues and we conclude that these Sen residues are in fact not directly involved in the binding of hPg to the Sen octamer. These residues are buried in the minor interface of Sen and are unavailable for this binding interaction. This confirms an earlier observation on the energetics of octamer formation (56). Hydrogen bonding between these residues and K^334^ and electrostatic interactions with E^335^ help to stabilize these residues in the Sen octamer as well as providing additional stability for the octamer as whole. Removal of these residues or altering them by mutagenesis decreases the stability of the octamer and result in dissociated ordered structures consisting of tetramers, hexamers, and heptamers as shown in **Figure 12**.

Since C-terminal K^435^ of Sen does not serve as a hPg binding site and since hPg binds through its LBS to Sen, hPg must be interacting through an isosteric lysine present on Sen. Lysine side chains, alone, would not provide the strength of interaction at any of the LBS of hPg. A side-chain carboxylate appropriately located must also serve to generate an internal lysine. We previously reported that a Lys side-chain and a Glu side-chain rigidly held in place at i, i+3 by one turn of an α-helix served as a strong binding site for hPg on another GAS PgR (57, 58). There are only five possibilities on a Sen protomer that present such a conformation, *viz*., K^191^-E^195^, E^220^-K^224^, K^224^-E^227^, E^281^-K^285^, K^304^-E^308^ (**Figure 11**). Thus, it is reasonable to propose that one or more of these sites interact with hPg. Additionally, as shown here, the internal potential lysine isostere encompassing K^252^ and K^255^ (34) is also not involved in this binding. These residues are present on L3 which is a very flexible loop without the rigidity to serve in this capacity. A flexible loop likely does not present the proper geometry to become a stable lysine isostere. While the high-resolution model of Sen developed herein supports the lack of interaction of hPg and Sen in solution, as well as the lack of involvement K^252,255^ and K^434,435^ in the binding of hPg to surface-adsorbed Sen. We stress that these conclusions apply to symmetrical octameric (and likely hexameric) enolases. In cases of mammalian enolases, which are dimeric, the minor interface does not exist and the enolases are free to interact with C-terminal lysine residues.

We propose that the differences in experimental data regarding Sen/hPg binding can be explained by subtleties in the experimental conditions used for different studies. It is clear that a specific perturbation of the Sen conformation, when bound to surfaces, is needed to allow access of hPg to its binding sites on Sen. In previous *in vitro* studies in which the binding of hPg to Sen mutants were studied, Sen was adsorbed to surfaces at pH values of 9.6 (27) and/or pH 4.0 (34), and the protein bound to the artificial surface is that which exists at these pH values. At pH 9.6, we find that the structures of the Sen mutants are perturbed (**Figure 2B**). Since C-terminal Lys residues at positions 434 and 435 stabilize the WT-Sen conformation, mutating these residues further destabilized the octameric structure **(Figure 2B)** and this type of perturbation of the octamer leads to a loss of hPg functional binding. It is interesting to note that when dissociated **(Figure 2B)**, only monomers are observed without any intermediate association states, demonstrating that Sen is in monomer-octamer equilibrium.

In conclusion, while C-terminal Lys residues of proteins are well known to be involved in interactions with hPg, and this is true for mammalian dimeric enolases, this is not the case for many prokaryotic enolases, many of which do not even possess Lys residues at this position. In the case of the closed octamer, Sen, the C-terminal lysine residues are present in the minor interface between two monomers and do not function as hPg binding loci. These minor interfaces are not present in mammalian dimeric enolases, which associate through major interfaces, and thus C-terminal lysine residues are available for hPg binding. The model of hPg binding to Sen (59) that best fits our ideas is that one molecule of hPg binds to the non-native, but fully associated, Sen octamer with a slightly relaxed octameric Sen conformation. The conformational changes in the octamer are subtle and may differ depending on the surfaces to which they are bound, but in the case of Sen, an octamer is required for functional binding of hPg.

## Data Availability

The cryo-EM coordinates of the model of octameric enolase have been deposited in the Worldwide Protein Database (PDB) database with the identifier 7UGU and the 2.6Å map has been deposited in the Electron Microscopy Protein Database (EMDB) with the identifier 26406. The cryo-EM coordinates of the model of octameric enolase [K^252,255,434,435^A] have been deposited in the Worldwide Protein Database (PDB) with the identifier 8DG4 and the 3.1Å map has been deposited in Electron Microscopy Protein Database (EMDB) with the identifier 27407.

## Acknowledgments

The authors thank Dr. Thomas Klose, technical Director of the Purdue University cryo-EM facility, for the expert training and assistance of the personnel involved in this study. All cryo-EM data were obtained at this facility.

## Author Contributions

ST-F., BR, and YAA, performed all experiments. FJC designed the study and wrote the manuscript.

