## Supplementary Data for "A high-resolution single-particle cryo-EM hydrated structure of *Streptococcus pyogenes* enolase offers insights into its function as a plasminogen receptor"

**Supplementary Table 1S. Primers employed to generate Sen variants**

| No. | Primer | Sequence* |
|---|---|---|
| P1 | SEN-WT-FP | 5’-GCCATATGTCAATTATTACTGATGTATAC-3’ |
| P2 | SEN-WT-RP | 5’-CCGGATCCCTATTTTTTTAAGTTATAGAATG -3’ |
| P3 | SEN-252,255-RP | 5’-CATCATCAGAATTTTACGACgcAGAACGTgcAGTT TACGACTAC-3’ |
| P4 | SEN-252,255-FP | 5’-GTAGTCGTAAACTgcACGTTCTgcGTCGTAAAATTCTGA TGATG-3’ |
| P5 | SEN-434,435-RP | 5’-CCGGATCCCTATgcTgcTAAGTTATAGAATGATTTGATA-3[illegible] |

* The noncapitalized nucleotides in the primers were mutagenic.

**Table 2S. Cryo-EM Parameters**

| | WT-Sen:<br>PDB-7UGU, EMDB-26406 | Sen[$K^{252,255,434,435}$A]<br>PDB-8DG4 EMD-27407 |
|---|---|---|
| Refinement Program | Cryosparc | Cryosparc |
| Magnification | 109,000 | 109,000 |
| Voltage, kV | 300 | 300 |
| Electron Exposure, e-/Å$^2$ | 61.37 | 61.37 |
| Defocus Range | -1.1 to -3.2 | -1.0 to -3.4 |
| Pixel Size, Å | 0.437 | 0.437 |
| Initial Particle images | 813,556 | 995,788 |
| Final Particle Images | 442,256 | 310,160 |
| Symmetry Imposed | D4 | D4 |
| Resolution unmasked,<br>FSC threshold 0.143Å | 3.2 | 3.9 |
| Resolution masked,<br>FSC threshold 0.143Å | 2.6 | 3.3 |
| FSC threshold | 0.143 | 0.143 |
| **Model Design** | | |
| Model resolution, Å | 3.5 | 3.5 |
| FSC threshold | 0.5 | 0.5 |
| Model resolution range | 5.4 | 3.5 |
| Map sharpening β factor (Å$^2$) | -21 | -0.78 |
| Refinement Program | Phenix, Chimera X | Phenix, Chimera X |
| Number of atoms, non-H | 26,392 | 26,264 |
| Protein Residues | 3,488 | 3,488 |
| Ligands | 0 | 0 |
| **B-factors (Å$^2$)** | | |
| Protein | 31 | 43 |
| **R.M.S Deviations** | | |
| Bond Length (Å) | 0.004 | 0.003 |
| Bond Angle (°) | 1.092 | 0.899 |
| **Validation** | | |
| MolProbity Score | 2.28 | 2.28 |
| Clash Score | 0.02 | 0 |
| Poor rotamers, % | 0.74 | 0.61 |
| Ramachandran favored, % | 94.9 | 96 |
| Ramachandran allowed, % | 5 | 4 |
| Ramachandran disallowed, % | 0.1 | 0:06 |
| EMringer score | 2.49 | 2.44 |

94.9% favored, 5% allowed, 0.1% outliers and an overall clash score at 0.02.

**Supplementary Figure 1S**

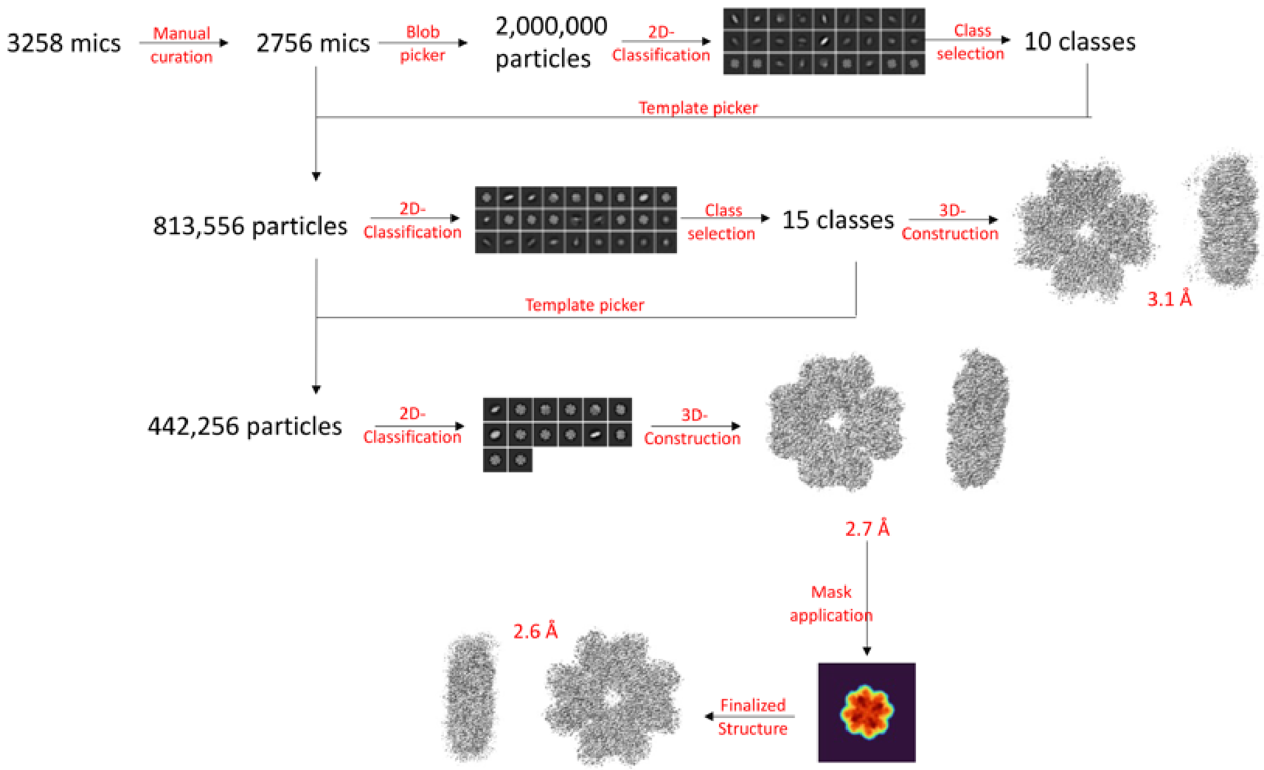


**Supplementary Figure 1S.** Workflow describing the procedure for generating the finalized map structure of WT-Sen in Cryosparc. The process of performing the methods is further described in the Experimental Results of this manuscript. The resolution of the maps was determined by 0.143 FSC.

**Supplementary Figure 2S**

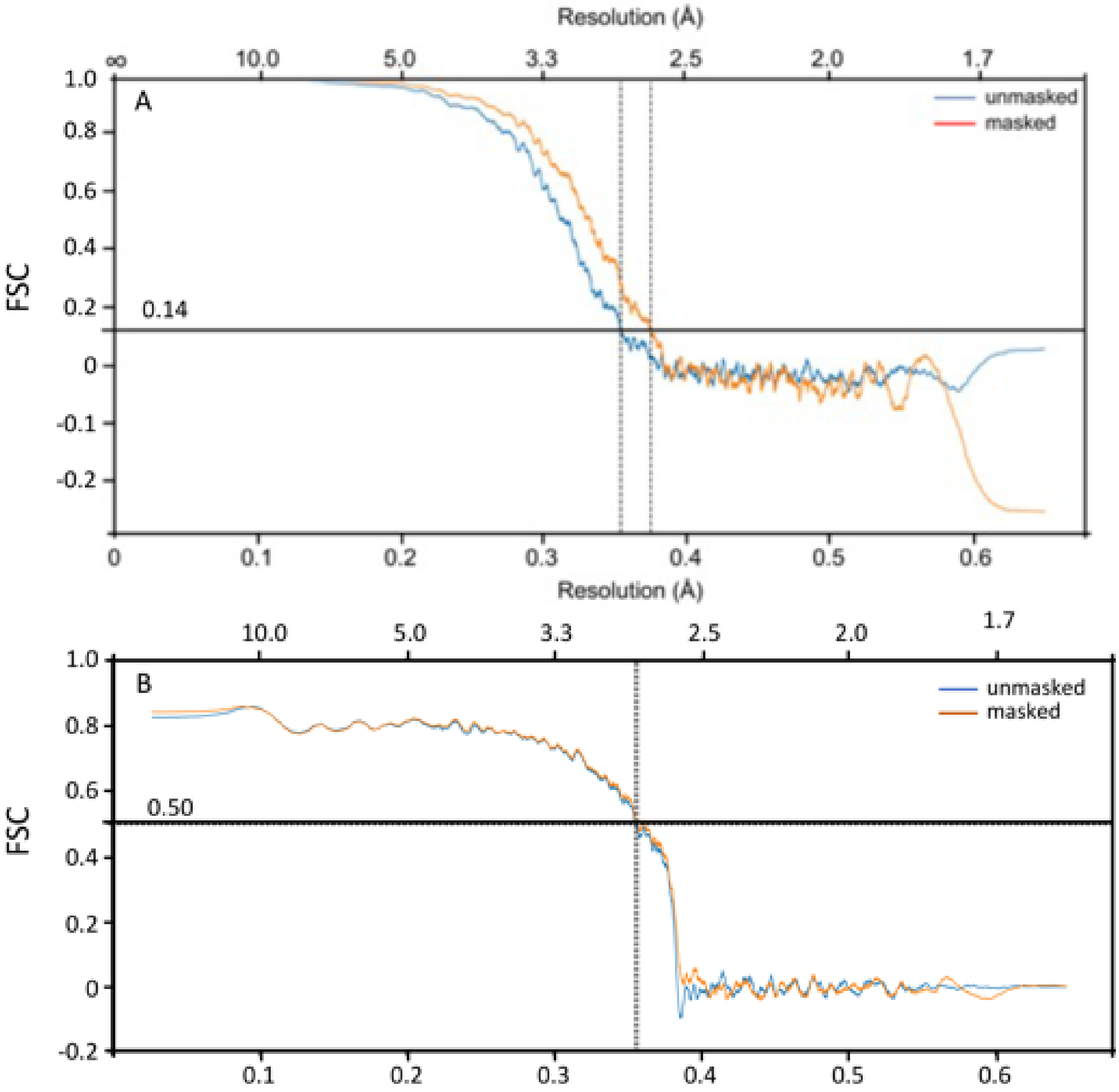


**Supplementary Figure 2S.** FSC plots used to determine the resolution of WT-Sen cryo-EM map and correlation between the map and the model. (A) The FSC plot at 0.143 threshold confirms the final resolution of the cryo-EM map to be 2.6Å. (B) The FSC plot at 0.5 threshold is used to compare the fit.
